# Phenolic Chemical Defense Contributes to Resistance against Multiple Soybean Cyst Nematode Populations in Wild Soybean

**DOI:** 10.64898/2026.09.12.751195

**Authors:** Hengyou Zhang, Neha Mittal, Xu Li, Zenglu Li, Chunying Li, Han-Yi Chen, Eric Davis, Changbao Li, Qijian Song, Yong-qiang Charles An, Bao-Hua Song

**Affiliations:** Department of Biological Sciences, University of North Carolina at Charlotte, Charlotte, NC 28223, USA; Northeast Institute of Geography and Agroecology, Key Laboratory of Soybean Molecular Design Breeding, Chinese Academy of Sciences, Harbin 150081, China. Thanks; Plant for Human Health Institute, North Carolina State University, Kannapolis, NC 28081, USA; Department of Plant and Microbial Biology, North Carolina State University, Raleigh, NC 27695, USA; Institute of Plant Breeding, Genetics and Genomics and Department of Crop and Soil Sciences, University of Georgia, GA 30602, USA; Department of Plant Pathology, North Carolina State University, Raleigh, NC 27607, USA; Syngenta Research Triangle Park, Durham, NC 27709, USA; United States Department of Agriculture, Agricultural Research Service, Soybean Genomics and Improvement Laboratory, Beltsville, MD 20705, USA; US Department of Agriculture, Agricultural Research Service, Midwest Area, Plant Genetics Research Unit, Donald Danforth Plant Science Center, St. Louis, MO, 63132, USA; School of Data Science, University of North Carolina at Charlotte, Charlotte, NC 28223, USA

**Keywords:** *Glycine soja*, soybean cyst nematode, *Heterodera glycines*, phenolic metabolism, chemical defense, metabolomics, transcriptomics, plant–nematode interaction

## Abstract

Soybean cyst nematode (*Heterodera glycines*, SCN) is one of the most damaging pathogens of soybean worldwide. Virulence diversity among SCN populations and reliance on a limited number of resistance sources present major challenges for durable SCN management. Here, we investigated a wild soybean (*Glycine soja*) genotype (WsR), that is resistant to two SCN populations, race 2 (HG type 1.2.5.7) and race 5 (HG type 2.5.7), and compared its transcriptional and metabolic responses with those of the susceptible genotype (WsS). WsR exhibited a substantially stronger defense-associated transcriptional response to both SCN populations, including preferential induction of genes associated with Ca²⁺/calmodulin and salicylic acid signaling. Integration of transcriptomic and time-resolved metabolomic analyses revealed convergence on phenolic metabolism, with enhanced accumulation of phenolic acids, flavonoids, and isoflavonoids in resistant WsR. Genes involved in phenolic biosynthesis and modification were also preferentially induced in WsR, linking transcriptional reprogramming with the observed metabolic response. Importantly, two resistance-associated phenolic compounds, 4-hydroxybenzaldehyde and 2,3-dihydroxybenzoic acid, directly increased mortality of SCN second-stage juveniles from both populations in a concentration-dependent manner. By connecting resistance-associated transcriptional responses and metabolic reprogramming with direct activity of specific phenolic compounds against SCN, our study provides functional evidence linking phenolic metabolism to chemical defense against SCN. Together, these findings identify enhanced phenolic chemical defense as a major component of resistance to multiple SCN populations in wild soybean and highlight wild soybean as a valuable source of molecular and biochemical diversity for SCN resistance.

## Introduction

Soybean (*Glycine max* L. Merr.) is an important legume crop grown worldwide that provides major sources of plant protein and vegetable oil (Carter et al., 2004). However, soybean production is severely challenged by soybean cyst nematode (SCN, Heterodera glycines Ichinohe); as one of the most damaging soybean pathogens, SCN causes substantial yield losses in the United States and other soybean-producing regions (Wrather and Koenning, 2006). SCN populations exhibit extensive virulence diversity and can shift in response to selection imposed by resistant cultivars. In the US, SCN has spread to all major soybean-growing states since its initial discovery in North Carolina in 1954 (Tylka and Marett, 2014). Resistance to one SCN population may not confer resistance to populations with different virulence profiles, creating a major challenge for durable SCN management. Often, an infested field comprises different virulence profiles, and a soybean cultivar resistant to one population might not be resistant to other or newly evolved populations. SCN diversity and race shifts pose major challenges for soybean scientists seeking effective SCN management strategies. To meet these challenges, identification of genetically diverse resistance sources effective against multiple SCN populations and uncovering their underlying mechanisms are equally important for guiding efforts to develop more durable SCN resistance. Currently, “HG type,” determined using seven soybean indicator lines, is increasingly used to describe SCN populations. For consistency with the historical designation of the populations used in this study, we refer to race 2 (HG type 1.2.5.7) and race 5 (HG type 2.5.7).

Thus far, studies on soybean–SCN interactions have been concentrated on a limited number of resistant soybean cultivars, particularly Peking and PI88788, which have contributed extensively to SCN resistance in commercial soybean cultivars in the United States. Studies have shown that resistance of Peking-type cultivars requires *rhg1*-a (Liu et al., 2017) and *Rhg4* (Liu et al., 2012), while resistance of PI88788-type cultivars is primarily associated with the *rhg1*-b allele (Cook et al., 2012), illustrating the genetic complexity of SCN resistance. The QTLs *rhg1* and *Rhg4*, conferring resistance to different SCN populations (HG types), have also been identified in other resistant soybean genotypes, demonstrating their central importance in soybean SCN resistance (Concibido et al., 2004, Vuong et al., 2010). On the other hand, extensive deployment of limited resistance sources, particularly PI88788, has increased genetic vulnerability as SCN populations capable of reproducing on these resistance sources have become more prevalent (Niblack et al., 2008). Thus, the identification of additional and genetically diverse sources of SCN resistance is needed.

Previous DNA marker-based and whole-genome sequencing studies have shown that wild soybean (*Glycine soja* Sieb. & Zucc.), the wild progenitor of cultivated soybean, retains a higher level of genetic diversity than cultivated G. max; substantial genetic variation present in wild soybean was lost during soybean domestication and improvement (Qi et al., 2014, Hyten et al., 2006). Previous studies have shown varying levels of resistance of *G. soja* to different SCN populations (HG types), and QTLs associated with SCN resistance have been identified in wild soybean (Kim et al., 2011, Zhang et al., 2017b, Zhang et al., 2016). Despite this genetic potential, the molecular and biochemical mechanisms contributing to SCN resistance in *G. soja* remain poorly understood.

Plants synthesize numerous natural products that are critical in plant defence against pathogens and herbivores (War et al., 2012). Phenolic compounds, including phenolic acids, flavonoids, and isoflavonoids, constitute an important component of plant chemical defence, and isoflavonoids are particularly important specialized metabolites in legumes. However, the contribution of phenolic chemical defence to SCN resistance remains incompletely understood. Much of the present knowledge of soybean responses to SCN has come from transcriptomic analyses or studies of individual defense-associated metabolites (Wan et al., 2015, Kandoth et al., 2011, Klink et al., 2007a, Klink et al., 2007b, Mazarei et al., 2011). Although these studies are insightful, an important gap remains between the transcriptional responses associated with multiple SCN resistance and the chemical defenses that may ultimately affect nematode establishment, development, or survival.

Untargeted metabolomics provides an opportunity to characterize these chemical responses at a broader scale and, when integrated with transcriptomic analysis, to connect infection-responsive genes with changes in defense-associated metabolites. Liquid chromatography–mass spectrometry (LC-MS) enables broad profiling of metabolites in complex biological samples (Huang and Barker, 1991, Nguyen et al., 2013). Given the great potential of *G. soja* to provide genetic and biochemical diversity for soybean improvement, integrating transcriptomic and metabolomic analyses provides a powerful approach for identifying molecular and chemical responses associated with SCN resistance in wild soybean.

In the present study, we investigated transcriptional and metabolic responses to two SCN populations, race 2 (HG type 1.2.5.7) and race 5 (HG type 2.5.7), in resistant and susceptible *G. soja* genotypes. We combined comparative RNA sequencing with time-resolved untargeted LC-MS metabolomics to address several related questions: 1) Does resistance to multiple SCN populations involve shared molecular and metabolic responses? 2) Which defense-associated pathways distinguish the resistant from the susceptible genotype? 3) Do transcriptomic and metabolomic responses converge on specific branches of secondary metabolism? 4) Most importantly, can metabolites associated with the resistant response directly affect SCN survival? By integrating molecular, metabolic, and functional analyses, we sought to bridge the gap between resistance-associated transcriptional responses and the chemical defenses that may contribute to SCN resistance in wild soybean.

## Results

### *Glycine soja* genotype WsR exhibits resistance to two races of SCN

The resistance responses of the two wild genotypes (WsR and WsS) to two races of SCN were determined by inoculating two-day-old seedlings with 2,500 fresh eggs. Thirty-five days after inoculation, female cysts were counted under a stereoscope. WsR showed resistance to both races (FI < 10%), while WsS was susceptible to both races (FI > 60%) (Fig. 1a).

**Fig. 1.**
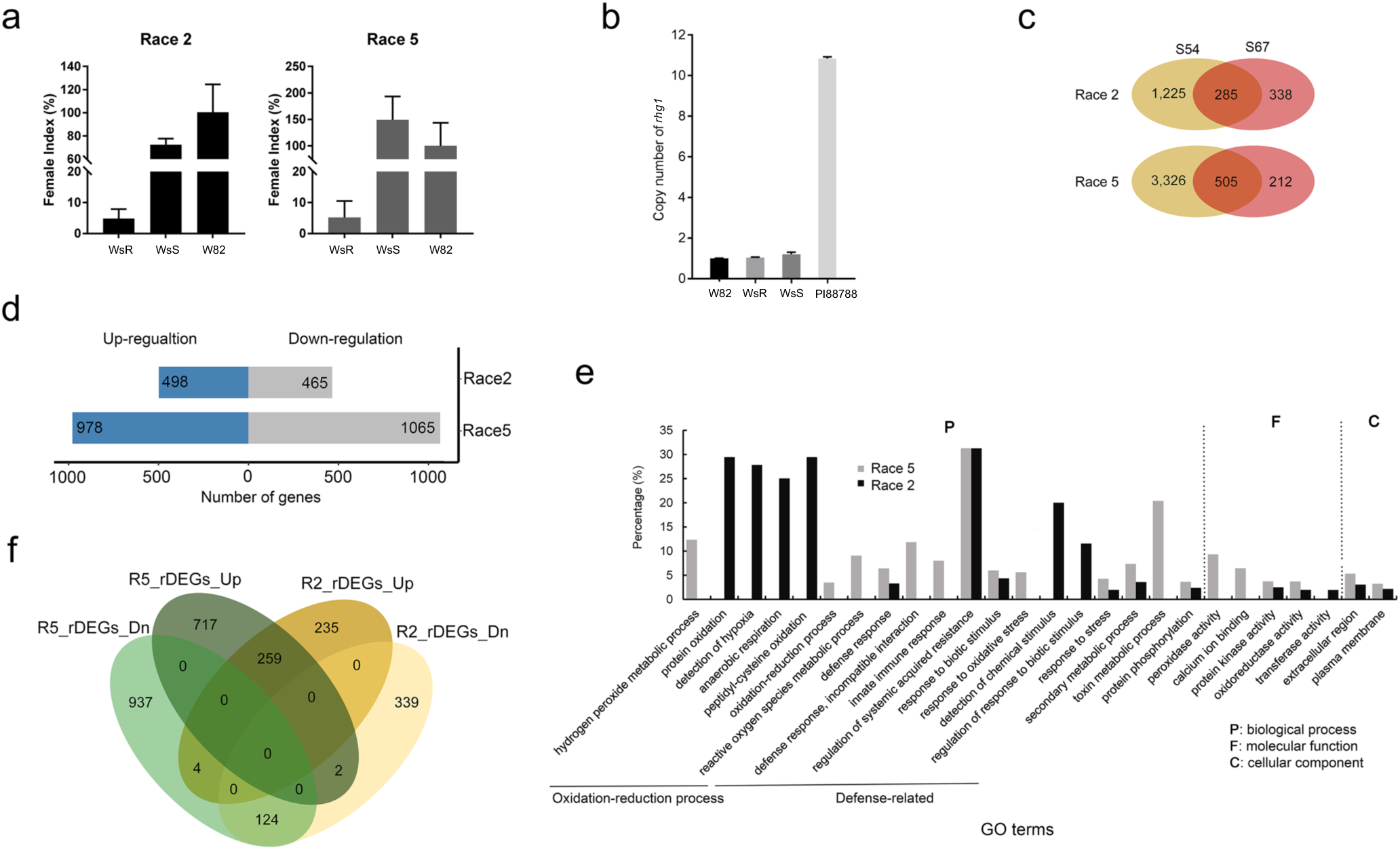
Phenotypic evaluation and global transcriptomic analysis of SCN responses in wild soybean. **a** Female indices of two *Glycine soja* genotypes, WsR and WsS, following inoculation with SCN races 2 and 5. *Glycine max* cv. Williams 82 was used as the susceptible check. **b** Copy number of Rhg1 in WsR and WsS compared with the SCN-susceptible soybean cultivar Williams 82 (W82) and the SCN-resistant accession PI 88788, which have known Rhg1 copy numbers. **c** Venn diagrams showing the numbers of differentially expressed genes (DEGs) between SCN-inoculated and corresponding control roots in WsR and WsS in response to races 2 and 5. **d** Numbers of up- and downregulated relative DEGs (rDEGs) identified for races 2 and 5. **e** GO enrichment analysis of upregulated rDEGs in response to races 2 and 5. **f** Venn diagram showing rDEGs shared between responses to races 2 and 5 and those specific to each race. **P < 0.01.

*Rhg1* and *Rhg4*, identified in soybean cultivars Peking and PI88788, are the two major SCN resistance loci underlying widely used resistance sources in cultivated soybean (Liu et al., 2017, Liu et al., 2012, Cook et al., 2012, Bent, 2022, Miraeiz et al., 2026a). *Rhg1* displays copy-number variation, with resistant *G. max* accessions typically carrying multiple copies of the locus (Cook et al., 2012). Genomic qPCR showed that WsR contained a single copy of *rhg1*, similar to the susceptible genotypes WsS and Williams 82 (Fig. 1b). In addition, we examined WsR using three molecular markers developed to distinguish resistance-associated genotypes at *rhg1* and *Rhg4* (Shi et al., 2015). WsR exhibited marker patterns similar to the susceptible genotype Lee74 and distinct from the resistant cultivars Peking and PI88788 (**Table 1**). Together, these results indicate that resistance in WsR differs from the *rhg1/Rhg4*-associated resistance represented by Peking and PI88788.

### WsR exhibits a stronger transcriptional response to *Heterodera glycines* infection than WsS

To identify transcriptional responses associated with SCN resistance in *G. soja*, we used RNA-seq to comparatively examine transcriptomic changes in WsR and WsS following infection with races 2 and 5. Root tissues collected at 3, 5, and 8 days post-inoculation (dpi) were pooled to maximally capture the transcriptome variation associated with the resistance response. In total, 24 libraries, 12 per race, were constructed for transcriptome sequencing (Table S1). RNA sequencing generated approximately 20 million reads per library. Over 80% of quality-controlled reads, on average, were uniquely mapped to the *Glycine max* reference genome.

Multidimensional scaling analysis showed that biological replicates clustered together and that experimental conditions were clearly separated (Fig. S1a). Differential expression analysis revealed that WsR exhibited a substantially stronger transcriptional response than WsS following infection by either race. Race 2 infection resulted in 1,510 and 612 differentially expressed genes (DEGs) in WsR and WsS, respectively, whereas race 5 infection resulted in 3,840 and 726 DEGs, respectively (Fig. 1c). Only a subset of the WsR-responsive genes was also differentially expressed in WsS. For race 2, 285 of the 1,510 WsR DEGs (18.9%) were shared with WsS, representing 46.6% of the WsS DEGs. For race 5, 514 of the 3,840 WsR DEGs (13.4%) were shared with WsS, representing 70.8% of the WsS DEGs (Fig. S1b). Thus, WsR mounted a substantially broader transcriptional response to SCN infection than the susceptible genotype WsS.

To identify infection-responsive genes showing stronger responses in WsR, we introduced the concept of relative DEGs (rDEGs). A DEG with a fold-change value in WsR that was >1.5-fold higher than that in WsS under infection by one race was designated a rDEG. This analysis identified 962 and 2,043 WsR-enriched responsive genes following race 2 and race 5 infection, respectively (Fig. 1d).

Hierarchical clustering separated these genes into two major groups comprising genes predominantly induced or suppressed in WsR (Fig. S2a). The majority of rDEGs identified in response to races 2 and 5 showed similar expression patterns, suggesting that WsR contains a group of genes potentially exhibiting shared responses to both SCN populations. GO enrichment analysis of the induced genes identified multiple defense-associated categories. Several terms, including regulation of systemic acquired resistance, response to biotic stimulus, response to stress, protein kinase activity, and plasma membrane, were enriched following infection by both races. In contrast, genes suppressed in WsR were enriched for functions associated with photosynthesis, transmembrane transport, chlorophyll binding, and cell-wall-related metabolism (Fig. S2b,c).

### Defense signaling pathways are preferentially activated in resistant WsR

To identify genes showing enhanced responses in WsR following infection by both races, we compared the two WsR-enriched gene sets. This analysis identified 383 shared genes, including 259 induced and 124 suppressed genes (Fig. 1f). GO and KEGG enrichment analyses showed that the induced genes were strongly associated with plant defense. “Regulation of systemic acquired resistance” was among the most significantly enriched GO terms (*q* < 0.05), while “plant–pathogen interaction” was among the enriched KEGG pathways (Fig. S2d). In contrast, the suppressed genes were predominantly associated with photosynthesis and metabolism.

Functional annotation of the commonly induced genes showed that most cDEGs belonged to gene families associated with plant disease defense (Table S3), including receptor-like kinases (RLKs), other protein kinases, NBS-LRR proteins, transcription factors, chitinases, laccases, and ankyrin-repeat proteins. Several well-characterized defense-associated kinases, including BAK1, BIR1, and SOBIR1, were strongly induced. Conversely, genes encoding FASCICLIN-like arabinogalactan proteins and cellulose synthases were among those suppressed, suggesting that cell-wall remodeling accompanies the WsR resistance response. Notably, multiple components associated with Ca²⁺/calmodulin (CaM) and salicylic acid (SA) signaling were strongly induced (Fig. 2; Table S2). Ca²⁺/CaM-associated genes included Ca²⁺-binding proteins, CaM-binding proteins, cyclic nucleotide-gated channels (CNGCs), and calreticulin 3 (CRT3), whereas SA-associated genes included *EDS1*, *SAMT1*, *GRX480*, *SARD1*, and *NIMIN1*. Network analysis further revealed extensive associations between these signaling components and other defense-related genes involved in transcriptional regulation, phosphorylation, immune recognition, hydrolysis, and ion transport (Fig. 2).

**Fig. 2.**
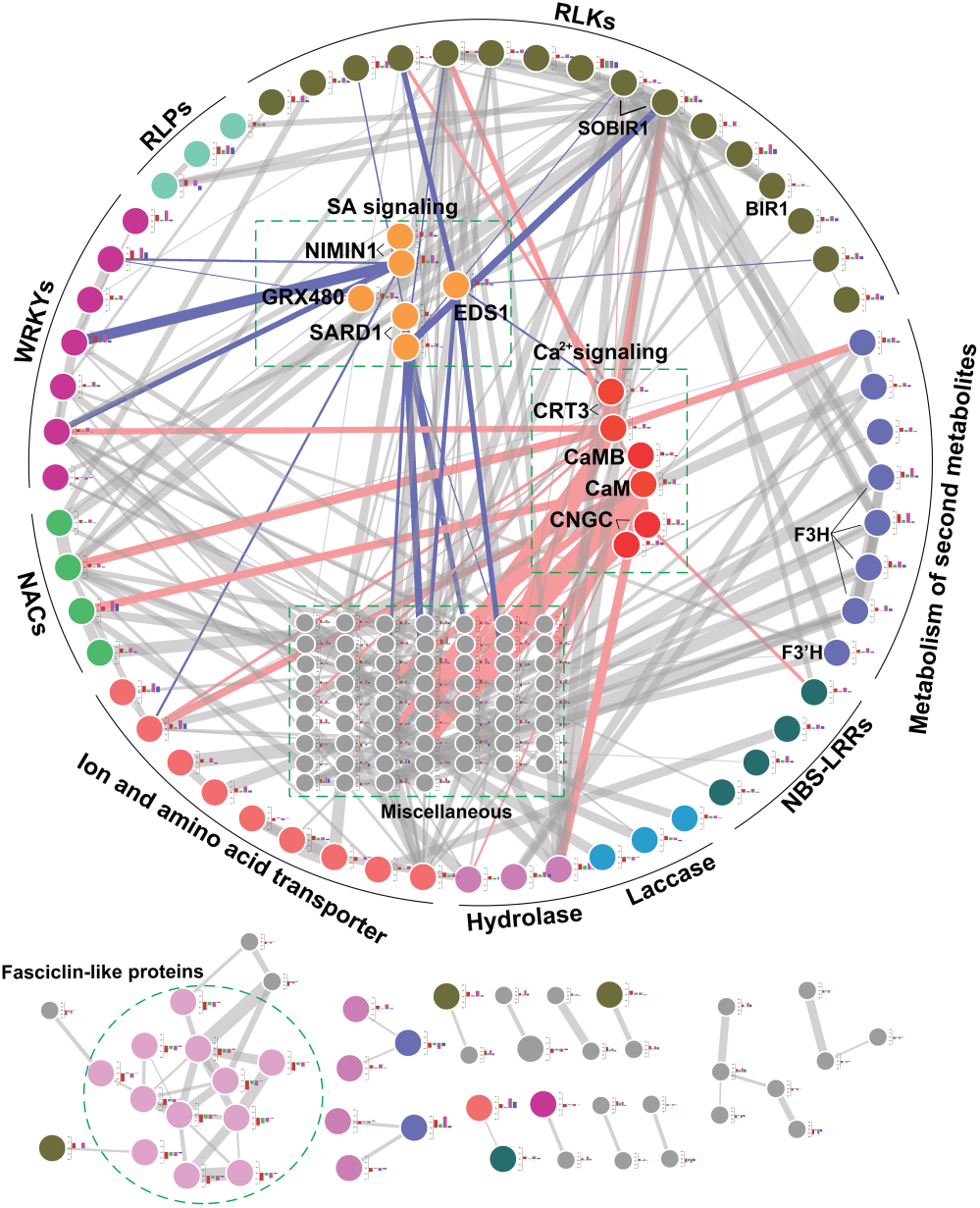
Interaction network of common-response differentially expressed genes (cDEGs). Each node represents a cDEG and is colour-coded according to its annotated gene family or functional category. Genes not assigned to the highlighted families are represented by small grey nodes. Edges connecting nodes represent predicted regulatory interactions, with edge weights corresponding to log-likelihood scores. Blue and orange edges highlight interactions connecting defence-related gene families with SA-signalling and Ca²⁺-signalling components, respectively. Histograms associated with individual nodes show RNA-Seq expression patterns across WsR_T_R2, WsR_C_R2, WsR_T_R5, and WsR_C_R5. RLKs, receptor-like kinases; RLPs, receptor-like proteins; NIMIN1, NIM1-INTERACTING 1; GRX480, SA-inducible glutaredoxin GRX480; SARD1, SYSTEMIC ACQUIRED RESISTANCE DEFICIENT 1; CRT3, calreticulin 3; CaM, calmodulin; CaMB, calmodulin-binding protein; CNGC, cyclic nucleotide-gated channel.

To examine the temporal responses of selected signaling genes, we quantified expression at 3, 5, and 8 dpi using qPCR. Five representative genes—a *CNGC*-like gene (*Glyma.03G257100*), *CaM* (*Glyma.06G258000*), *EDS1* (*Glyma.06G187300*), *PR1* (*Glyma.15G062400*), and *NIMIN1* (*Glyma.10G010100*)—were significantly induced in WsR following infection by both races, although their expression maxima occurred at different time points (Fig. 3). In contrast, these genes showed little or no induction in infected WsS roots. Two additional defense-signaling genes, *SAMT1* and *CAMTA*, similarly showed stronger induction in WsR than in WsS (Fig. S3a). Together, these results indicate preferential activation of Ca²⁺/CaM- and SA-associated defense signaling in resistant WsR.

**Fig. 3.**
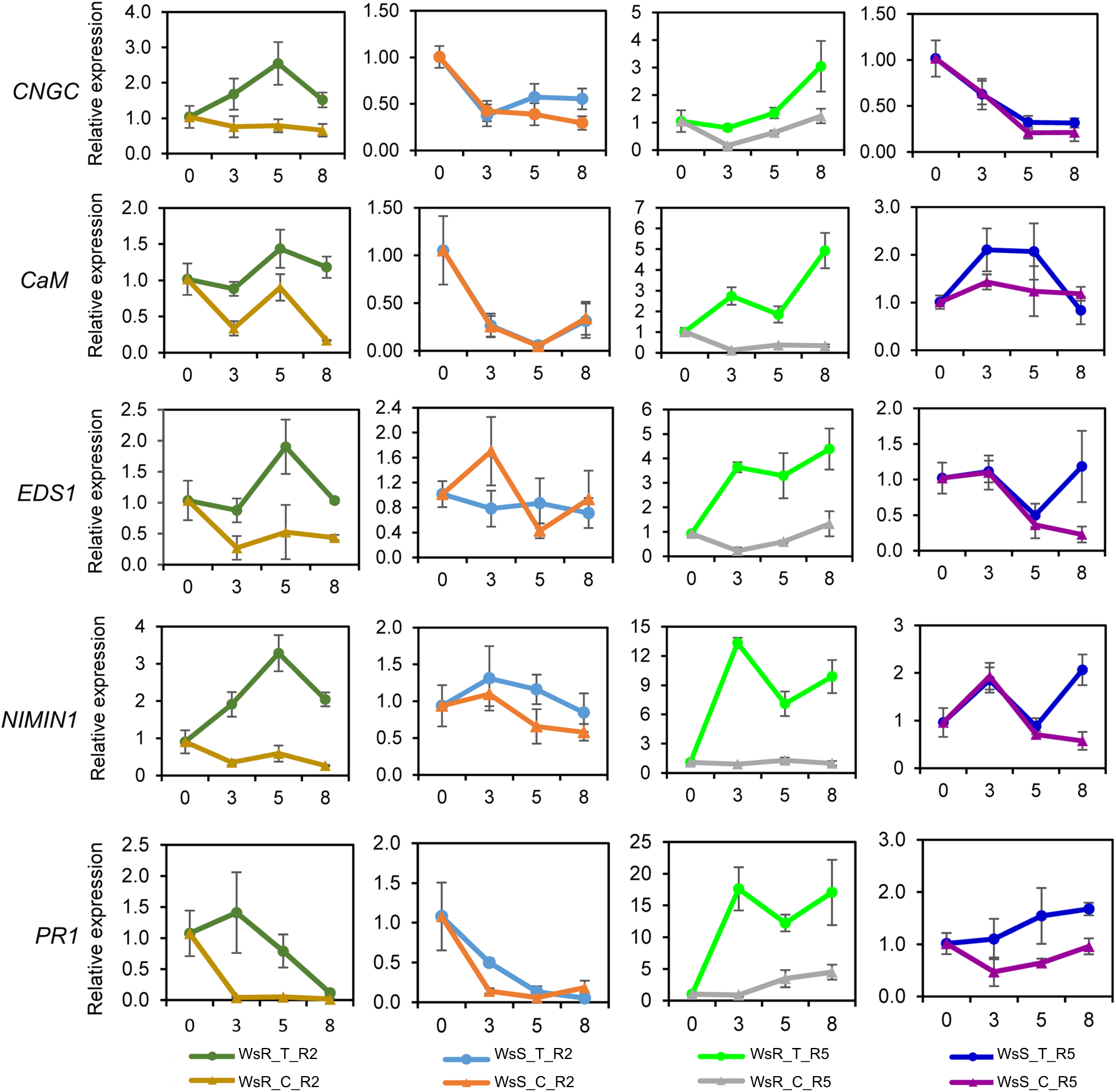
Time-course qPCR analysis of selected DEGs. Time-course expression analysis by qPCR of two Ca²⁺-related and three SA-signalling-related genes in roots of WsR and WsS following inoculation with SCN races 2 or 5, compared with the corresponding non-inoculated controls at 0, 3, 5, and 8 dpi. WsR_T_R2 and WsS_T_R2 denote race 2-inoculated roots; WsR_C_R2 and WsS_C_R2 denote the corresponding non-inoculated controls. WsR_T_R5 and WsS_T_R5 denote race 5-inoculated roots; WsR_C_R5 and WsS_C_R5 denote the corresponding non-inoculated controls.

### SCN infection induces extensive metabolic reprogramming in WsR

We next asked whether the resistance-associated transcriptional response was accompanied by changes in root metabolism. Untargeted LC-MS metabolomics detected 572 and 532 high-quality metabolite features in experiments involving races 2 and 5, respectively. Principal component analysis (PCA) separated samples according to both genotype and SCN treatment (Fig. 4a,b). Biological replicates clustered closely within each genotype, treatment, and time point. SCN infection induced extensive metabolic changes in WsR. A total of 71 metabolites were significantly increased following race 2 infection and 34 following race 5 infection, with 17 metabolite features increased in response to both races (Fig. 4c).

**Fig. 4.**
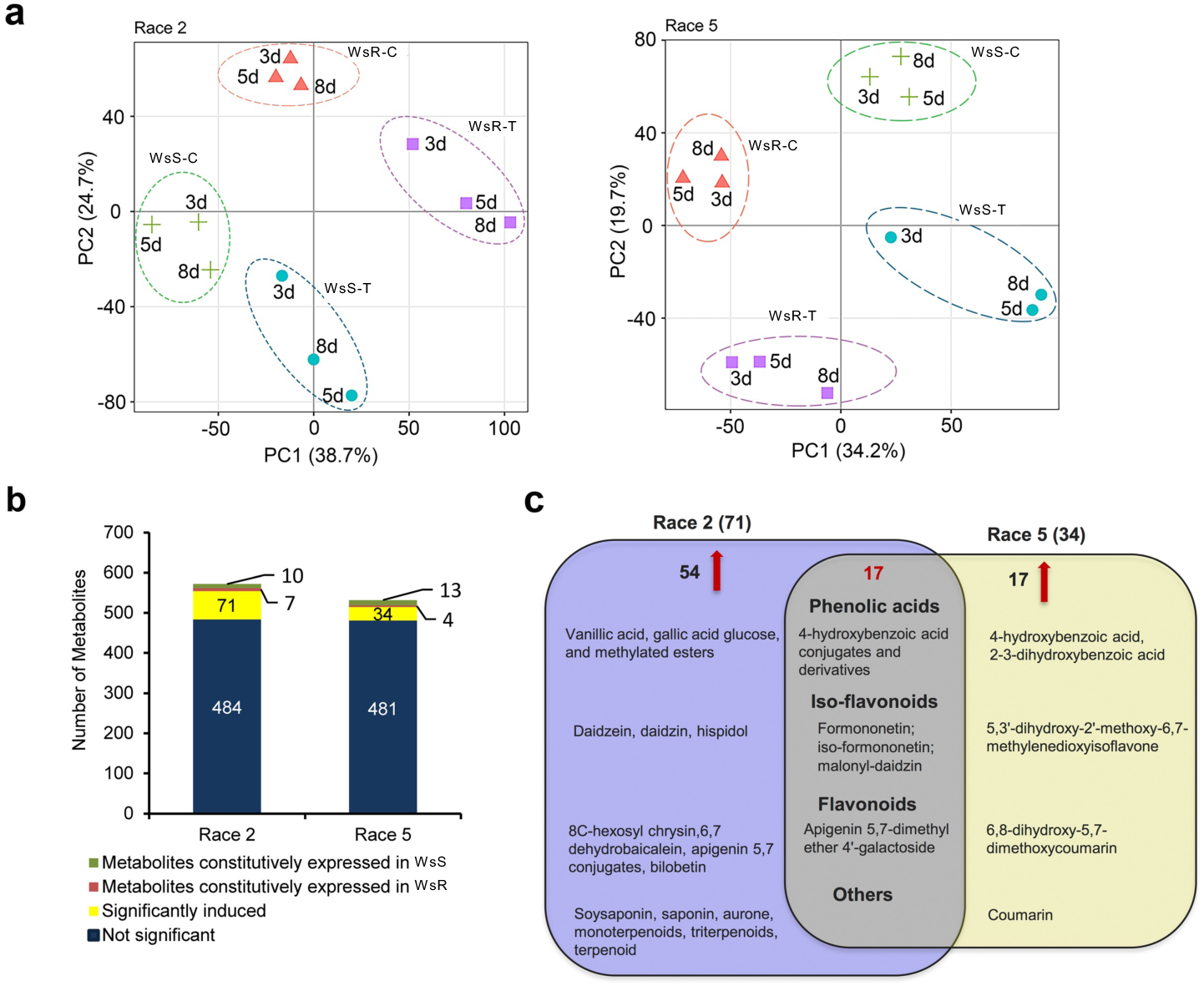
Global metabolomic profiles of wild soybean responses to SCN races 2 and 5. **a** Principal component analysis (PCA) of metabolite profiles from WsR and WsS roots following inoculation with SCN races 2 and 5 and their corresponding controls. Biological replicates cluster by experimental condition and are separated according to genotype and treatment. **b** Numbers of metabolites detected in roots inoculated with races 2 and 5, including constitutively different metabolites and SCN-induced metabolites. Constitutively different metabolites were defined as metabolites showing similar abundance between inoculated and control roots within one genotype but significantly different abundance between genotypes. **c** Venn diagram showing metabolites commonly induced by both races and those specifically induced by race 2 or race 5.

### Phenolic metabolism is strongly activated during the WsR response

Phenolic compounds constituted a major fraction of the annotated infection-responsive metabolites. Of the 47 annotated compounds increased following race 2 infection, 26 (55.3%) were phenolic compounds, including phenolic acids, isoflavonoids, and flavonoids. Similarly, 13 of 22 annotated compounds (59.1%) increased following race 5 infection were phenolics (Table S4). Terpenoids and saponins were also represented among the infection-responsive metabolites. Comparison of the two metabolomics datasets identified 12 annotated metabolites that increased following infection by both races, nine of which (75%) were phenolic compounds (Fig. 4c; Table S4).

To examine relationships among the infection-responsive metabolites, we mapped the shared metabolites together with selected race-specific compounds, including daidzein and daidzin, onto phenolic biosynthetic pathways (Fig. 5). The mapped metabolites represented phenolic-acid, isoflavonoid, and flavonoid branches derived from phenylalanine-associated metabolism. The resistance-associated phenolic metabolites also exhibited dynamic accumulation during infection. Under race 2 infection, many increased progressively beginning at 3 dpi, whereas under race 5 infection, several showed pronounced accumulation at 8 dpi. For both races, most reached their highest abundance at 8 dpi (Fig. 5). Some of these metabolites also increased in infected WsS roots, but their accumulation was generally weaker than in WsR. Together, these results identify enhanced phenolic metabolism as a major component of the WsR response to SCN infection.

**Fig. 5.**
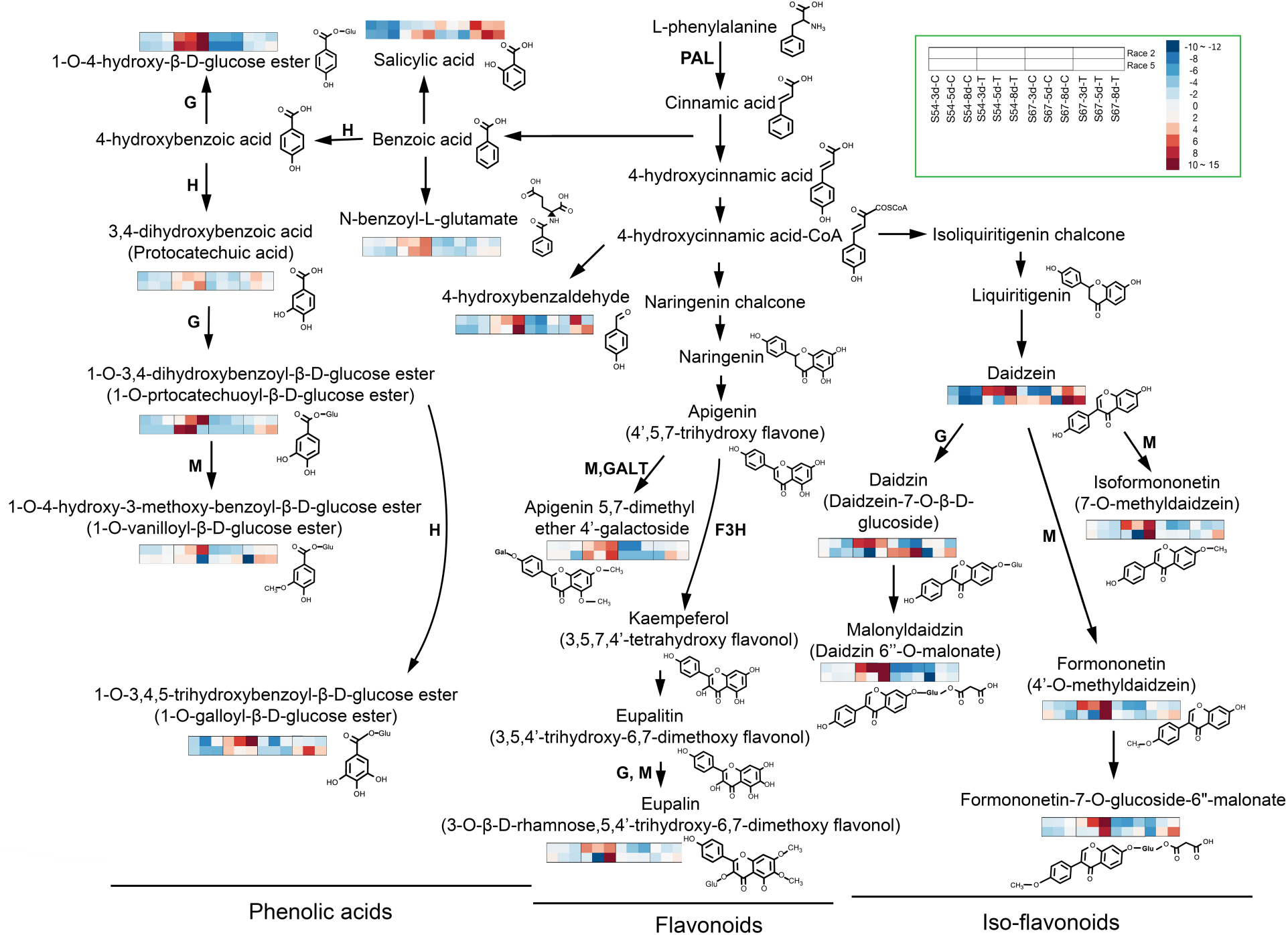
Phenolic biosynthesis pathways associated with SCN responses in wild soybean. Proposed metabolic relationships among constitutively abundant and SCN-induced metabolites identified in WsR roots, together with their chemical structures and positions within phenolic biosynthetic pathways. Heat maps show metabolite abundance across the treatment time course. Each square represents one biological replicate, and color intensity indicates relative metabolite abundance according to the scale shown. Sample order is indicated in the figure. Putative enzymatic steps involving hydroxylases, methyltransferases, glycosyltransferases, and galactosyltransferases are indicated by H, M, G, and GALT, respectively. F3H, flavonoid 3-hydroxylase; PAL, phenylalanine ammonia-lyase.

### Selected phenolic acids exhibit direct nematicidal activity against SCN

To determine whether resistance-associated phenolic metabolites could directly affect SCN, we tested two water-soluble phenolic compounds identified in the metabolomics analysis, 4-hydroxybenzaldehyde and 2,3-dihydroxybenzoic acid, against freshly hatched second-stage juveniles (J2) of races 2 and 5. Both compounds significantly increased J2 mortality relative to untreated controls, and mortality increased with compound concentration (Fig. 6). Overall, both 4-hydroxybenzaldehyde and 2,3-dihydroxybenzoic acid exhibited nematicidal activity against races 2 and 5. These results provide direct evidence that selected phenolic compounds associated with the WsR metabolic response can adversely affect SCN.

**Fig. 6.**
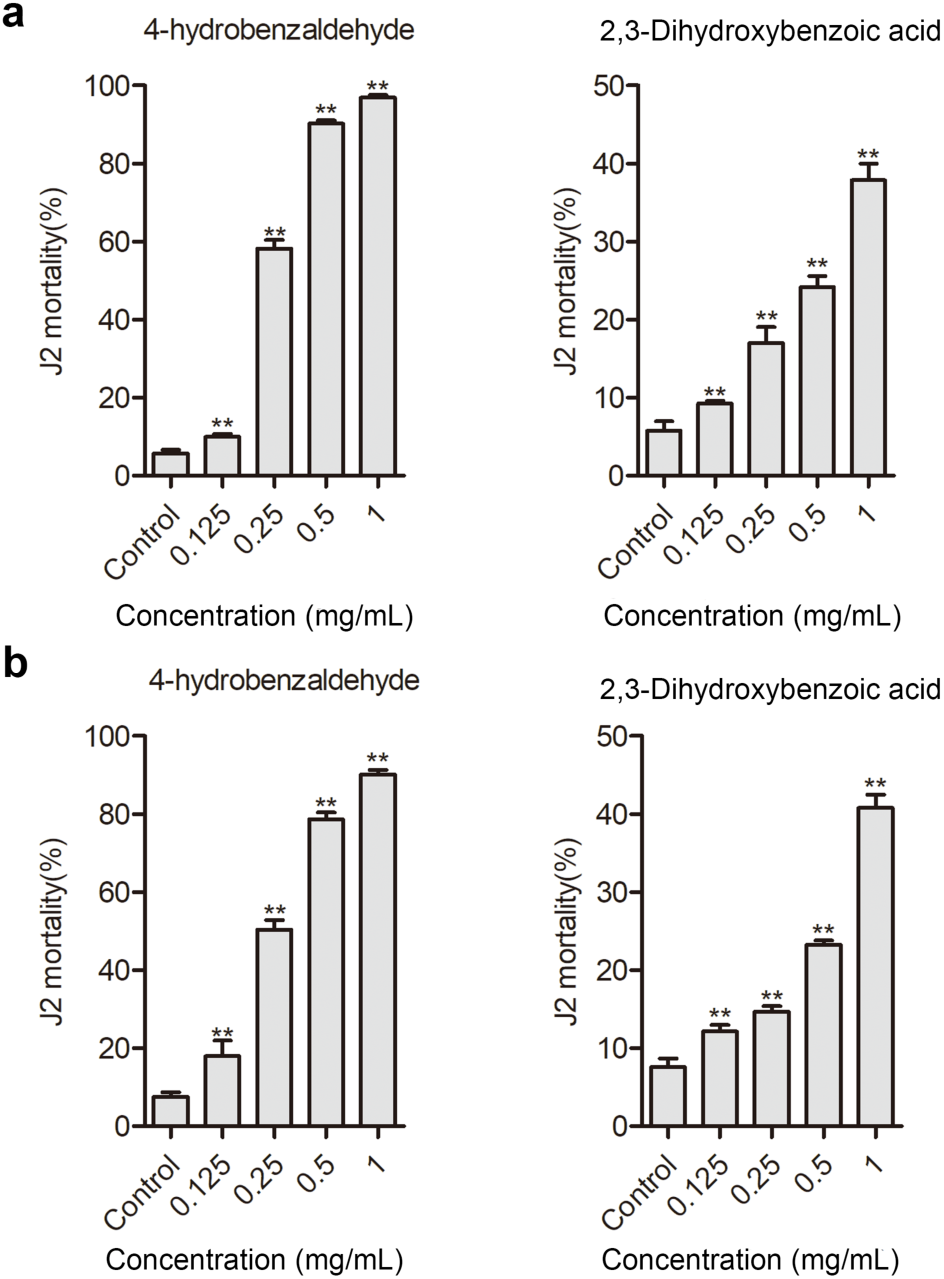
Nematicidal activity of two phenolic compounds against SCN second-stage juveniles (J2s) in vitro. Nematicidal activity of 4-hydroxybenzaldehyde and 2,3-dihydroxybenzoic acid against SCN J2s at the indicated concentrations. The x-axis indicates compound concentration (mg/mL), and the y-axis indicates J2 mortality (%). **P < 0.01 compared with the control.

### Transcriptomic and metabolomic analyses converge on phenolic biosynthesis

We next asked whether transcriptional changes in WsR were consistent with the enhanced accumulation of phenolic metabolites. Several genes encoding enzymes involved in phenolic, flavonoid, and isoflavonoid metabolism were induced following infection by both races (Fig. 7a). These included genes encoding flavanone 3-dioxygenases (*F3H*), flavonoid 3ʹ-hydroxylase (*F3ʹH*), O-methyltransferases (*OMT*), UDP-glycosyltransferases (*UGT*), galactosyltransferases (*GALT*), and glycosyl hydrolases.

**Fig. 7.**
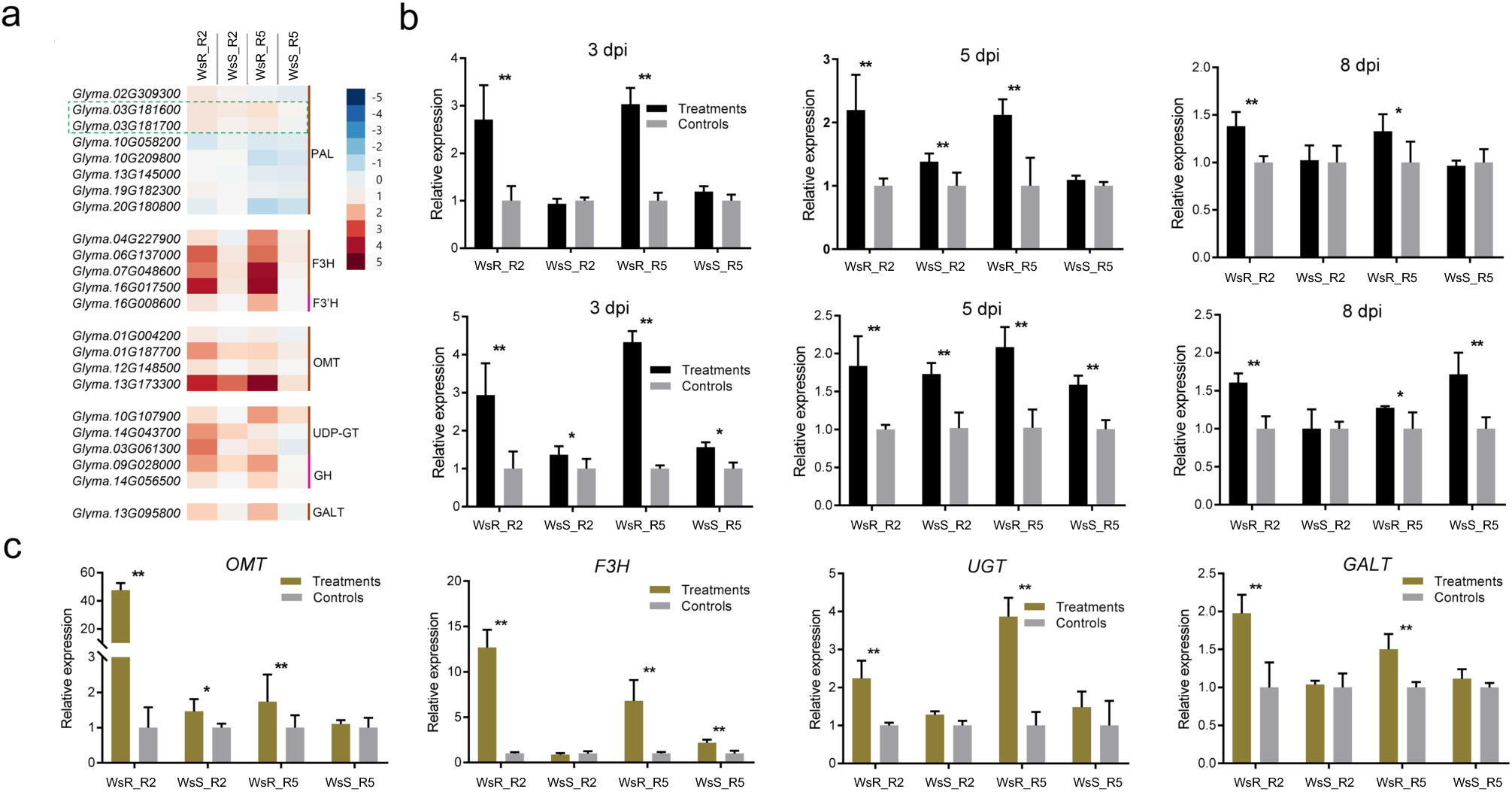
Expression patterns of candidate cDEGs associated with phenolic biosynthesis. **a** RNA-Seq expression patterns of candidate cDEGs associated with phenolic biosynthetic pathways. Colour intensity represents log₂ fold change (SCN-inoculated/control) according to the scale shown. **b** Expression analysis of two PAL genes highlighted in the dashed green box in a. Expression patterns of Glyma.03G181600 and Glyma.03G181700 are shown in the upper and lower panels, respectively. **c** Expression analysis of four selected candidate genes from a at 8 dpi. PAL, phenylalanine ammonia-lyase; F3H, flavonoid 3-hydroxylase; F3′H, flavonoid 3′-hydroxylase; OMT, O-methyltransferase; UGT, UDP-glycosyltransferase; GH, glycoside hydrolase; GALT, galactosyltransferase. *P < 0.05; **P < 0.01.

We subsequently used qPCR to examine selected pathway genes. Two *PAL* genes encoding phenylalanine ammonia-lyases, which catalyze an early step in phenylpropanoid metabolism, were also included. The selected pathway genes showed substantially stronger induction in infected WsR roots than in WsS (Fig. 7b,c). The two *PAL* genes exhibited particularly strong induction at 3 and 5 dpi, preceding the maximal accumulation of many phenolic metabolites observed at 8 dpi. Thus, independent transcriptomic and metabolomic analyses converge on enhanced phenolic metabolism in resistant WsR, supporting a model in which transcriptional activation of phenolic biosynthetic and modification pathways contributes to the accumulation of defense-associated metabolites during SCN infection.

## Discussion

### WsR represents an SCN resistance source distinct from *rhg1/Rhg4*-associated resistance

Wild soybean provides a valuable genetic resource for identifying SCN resistance mechanisms that may be underrepresented in cultivated soybean germplasm (Zhang et al., 2017c, Kofsky et al., 2018, Bent, 2022, Miraeiz et al., 2026a). The progenitor of *G. max*, *G. soja*, has a broad ecological and geographical distribution across East Asia and contains extensive genetic diversity that was reduced during soybean domestication and improvement (Zhou et al., 2015, Qi et al., 2014, Zhang et al., 2017a, Kofsky et al., 2018).

For SCN resistance, *rhg1* and *Rhg4* are the two major loci that have been most extensively characterized and deployed in cultivated soybean (Liu et al., 2017, Liu et al., 2012, Cook et al., 2012, Bent, 2022, Miraeiz et al., 2026a). Several observations indicate that WsR differs from the resistance sources represented by Peking and PI88788. First, WsR showed molecular-marker patterns at *rhg1* and *Rhg4* similar to those of the susceptible genotype Lee74 and distinct from Peking and PI88788 (Table 1). Second, WsR contained a single copy of *rhg1*, similar to susceptible WsS and Williams 82, rather than the elevated copy numbers characteristic of major *rhg1*-mediated resistance sources. Third, the three genes within the *rhg1* locus whose increased expression is associated with elevated *rhg1* copy number did not show differential expression between WsR and WsS or strong induction following SCN infection in our previous analysis (Zhang et al., 2017c).

Although these observations do not establish that WsR resistance is genetically independent of *rhg1* or *Rhg4*, they strongly suggest that WsR represents a resistance source distinct from the canonical *rhg1/Rhg4* configurations widely deployed in cultivated soybean. Identifying the genetic and biochemical components underlying this resistance may therefore expand the diversity of mechanisms available for improving SCN resistance in soybean.

### Resistance of WsR to multiple SCN races is associated with a distinct defense response

The extensive virulence diversity of *H. glycines* and shifts in field populations can reduce the durability of resistance derived from a limited number of genetic sources (Concibido et al., 2004, Niblack et al., 2008, Mitchum, 2016, Goverse and Mitchum, 2022, Miraeiz et al., 2026a). Identification of additional resistance mechanisms is therefore important for diversifying SCN resistance in soybean.

WsR showed strong resistance to both races 2 and 5, whereas WsS was susceptible to both. Resistance to two distinct SCN races does not establish universal or broad-spectrum resistance, but it demonstrates that the WsR phenotype is effective against multiple SCN populations with different virulence profiles.

The distinct molecular and metabolic responses observed in WsR further suggest that wild soybean may contain defense components complementary to those represented by commonly deployed cultivated resistance sources.

Rather than simply producing a larger number of infection-responsive genes, resistant WsR exhibited a qualitatively distinct defense response compared with susceptible WsS. Genes showing stronger responses in WsR were enriched for functions associated with immune signaling, including receptor-like kinases, transcription factors, Ca²⁺/CaM signaling components, and SA-associated regulators. In contrast, many of these genes showed weak or little induction in WsS.

Ca²⁺ signaling and SA-mediated defense are well-established components of plant immunity, and extensive interactions occur between these signaling systems (Lecourieux et al., 2006, Spoel and Dong, 2012, Miraeiz et al., 2026b). In WsR, genes encoding CNGCs, CaM-associated proteins, CRT3, EDS1, SARD1, NIMIN1, and other defense regulators were preferentially induced following infection. The time-resolved qPCR analysis further showed that representative genes from these pathways were activated at different stages of the early infection response.

These results are consistent with activation of Ca²⁺/CaM- and SA-associated defense signaling in WsR. However, because changes in Ca²⁺ flux and pathway activity were not measured directly, the transcriptomic data should be interpreted as evidence for activation of genes associated with these pathways rather than demonstration of a specific Ca²⁺–SA signaling cascade. Importantly, the transcriptional response was accompanied by substantial changes in secondary metabolism, providing a potential biochemical output of the enhanced defense response in WsR.

### Phenolic metabolism is a major component of the WsR response to SCN

The strongest convergence between the transcriptomic and metabolomic datasets occurred in phenolic metabolism. More than half of the annotated metabolites induced in WsR following infection by either race were phenolic compounds, and phenolics accounted for most of the annotated metabolites increased in response to both races. These metabolites represented several related branches of phenylalanine-derived metabolism, including phenolic acids, flavonoids, and isoflavonoids.

At the transcriptional level, genes encoding enzymes associated with these pathways, including *PAL*, *F3H*, *F3ʹH*, *OMT*, *UGT*, and *GALT*, were preferentially induced in resistant WsR. Recent functional studies provide additional support for the involvement of phenylpropanoid and isoflavonoid metabolism in SCN resistance. Overexpression of GmPAL genes enhanced SCN resistance and inhibited nematode development, while the isoflavones daidzein and genistein also suppressed SCN development (Yang et al., 2024). More recently, GmUGT88A1 was shown to regulate both SCN resistance and isoflavone accumulation, providing genetic evidence linking modification of isoflavonoid metabolism with SCN resistance (Jiang et al., 2025). The temporal relationship between gene expression and metabolite accumulation was also informative: strong induction of *PAL* genes at earlier time points preceded the maximal accumulation of many phenolic metabolites at 8 dpi.

Together, these observations support coordinated transcriptional and metabolic reprogramming of phenolic biosynthesis during the WsR resistance response. Modification enzymes such as methyltransferases, hydroxylases, and glycosyltransferases may further contribute to the structural diversity of the accumulated compounds through methylation, hydroxylation, and glycosylation.

Because many phenolic compounds function in plant defense, their preferential accumulation in WsR suggested that chemical defense could contribute to the observed SCN resistance. This hypothesis was directly examined using selected phenolic compounds identified in the metabolomics analysis.

### Phenolic compounds provide a functional link between metabolic reprogramming and SCN resistance

Plant phenolics are widespread secondary metabolites with diverse roles in plant defense (Saini et al., 2024). Flavonoids and isoflavonoids, including daidzein and formononetin, have previously been associated with defense responses in legumes (Yang et al., 2024). Phenolic acids, although less extensively studied in plant–nematode interactions, can also exhibit biological activity against plant pathogens and herbivores (Nguyen et al., 2013).

A particularly important result of this study is that two phenolic compounds associated with the WsR metabolic response, 2,3-dihydroxybenzoic acid and 4-hydroxybenzaldehyde, directly increased SCN J2 mortality in a concentration-dependent manner. Both compounds were active against races 2 and 5. This functional assay moves the association between phenolic accumulation and resistance beyond transcriptomic or metabolomic correlation and demonstrates that at least some components of the infection-induced phenolic profile can directly affect SCN. Importantly, the nematicidal activities demonstrated here should not be generalized to all phenolic compounds identified in WsR. Rather, the results establish direct activity for the two experimentally tested compounds and motivate further investigation of other resistance-associated metabolites.

### A proposed model for SCN resistance in wild soybean

Collectively, our results support a working model in which resistance of WsR to multiple SCN populations involves interconnected defense-associated signaling, immune responses, and phenolic chemical defense (Fig. 8). Following SCN infection, resistant WsR showed preferential induction of genes associated with Ca²⁺/CaM and SA signaling, together with enhanced expression of diverse immune- and defense-associated genes, including protein kinases, NBS-LRR proteins, and transcriptional regulators. In parallel, genes involved in phenolic biosynthesis and modification, including F3Hs, OMTs, and UGTs, were preferentially induced, accompanied by substantial accumulation of phenolic acids, flavonoids, and isoflavonoids. Importantly, direct nematicidal activity of 2,3-dihydroxybenzoic acid and 4-hydroxybenzaldehyde provides functional evidence linking this metabolic response to chemical defense against SCN. We therefore propose that enhanced phenolic chemical defense represents a major component of the WsR resistance response, operating within a broader network of defense-associated responses. Further genetic and biochemical analyses will be required to establish the causal relationships among these signaling, transcriptional, and metabolic components and to determine the genetic basis of resistance in WsR. This study therefore highlights two complementary opportunities provided by wild germplasm: identification of resistance mechanisms that differ from those dominating cultivated soybean and discovery of defense-associated metabolites and pathways that can inform future genetic improvement. Further genetic dissection of WsR will be required to determine the loci controlling this phenotype and to establish how the signaling and metabolic responses identified here are connected in planta.

**Fig. 8.**
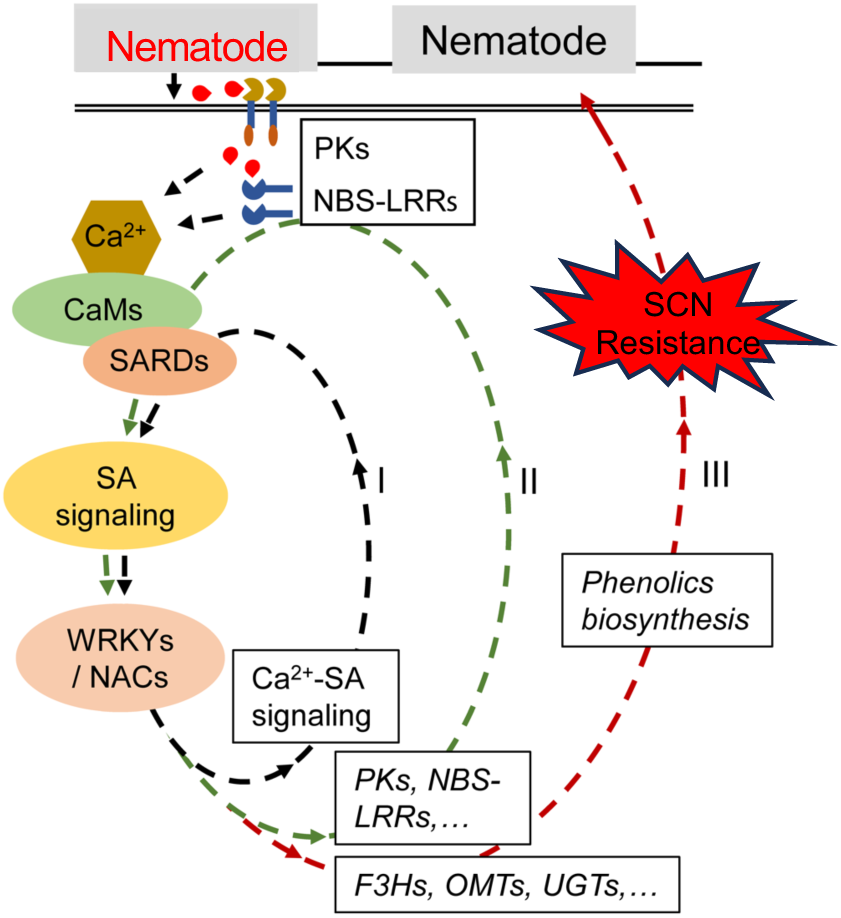
Proposed model of resistance to multiple SCN populations in wild soybean. Proposed model integrating transcriptional and metabolomic responses associated with resistance to multiple SCN populations in WsR. Following recognition of SCN (*Heterodera glycines*), receptor-like protein kinases (PKs) and NBS-LRR proteins are proposed to activate downstream defence signalling involving Ca²⁺, SA, WRKYs, and NACs. Enhanced defence signalling is associated with increased expression of genes encoding enzymes involved in phenolic biosynthesis, including F3Hs, OMTs, and UGTs, and with accumulation of phenolic acids, flavonoids, and isoflavonoids. Two identified phenolic compounds, 4-hydroxybenzaldehyde and 2,3-dihydroxybenzoic acid, exhibited nematicidal activity in vitro and may contribute to the resistant phenotype.

## Methods

### Plant materials and SCNs

Seeds of two *Glycine soja* genotypes, WsR (PI424093) and WsS (PI468396B), and seven *Glycine max* indicator lines (Peking, PI88788, PI90763, PI437654, PI209332, PI89772, and PI548316) for determining SCN HG types were obtained from the USDA Soybean Germplasm Collection (www.ars-grin.gov/). Williams 82 was used as a susceptible check in resistance screening. Two HG types of SCN, namely, 1.2.5.7 (known as race 2) and 2.5.7 (known as race 5), were used in this study. These two races were confirmed using the seven soybean indicators mentioned above with the methods previously described (Niblack et al., 2002). Both HG types were separately reared on soybean cv. Williams 82 grown in clay pots filled with sand under controlled greenhouse conditions (27 °C, 16 h light/8 h dark) for more than 30 generations. We previously reported the resistance responses of *G. soja* to SCN race 5 (Zhang et al., 2016). Here, together with examining *G. soja* resistance to SCN race 2, we re-evaluated resistance to race 5 and obtained consistent results.

The copy number of *rhg1* was quantified using genomic qPCR as described in Lee et al. (2015) (Lee et al., 2015). Determination of the genotype patterns of *rhg1* and *Rhg4* was performed as described by Shi et al. (2015) (Shi et al., 2015). The transcriptomic data of WsR and WsS responses to SCN race 5 were reported in our previous study (Zhang et al., 2016), and their responses to race 2 were investigated in the current study. Both data sets were analyzed using the same pipeline to identify shared infection-responsive genes.

### Plant preparation, SCN inoculation, and sample collection

Seed preparation, germination, transplanting, and SCN inoculation were performed as previously described (Zhang et al., 2017c). Briefly, *G. soja* seeds were surface sterilized with 0.5% sodium hypochlorite for 60 seconds, rinsed with autoclaved water, and then placed on wet sterile filter paper in a Petri dish for germination. After 2–3 days, each healthy seedling was transplanted into a container (Greenhouse Megastore, Danville, IL, USA) filled with sterile sand. Three days after transplantation, healthy seedlings were used for inoculation for resistance determination and sample collection for RNA-seq. A randomized complete block design was used for the seedling arrangement.

The resistance response of *G. soja* to SCN was determined as previously described (Zhang et al., 2016). The roots of each individual plant were inoculated with 2,500 fresh SCN eggs, and female cysts were collected from individual roots and the soil and counted under a stereomicroscope 35 days after inoculation. Four biological replicates per genotype were used, and the average number of females was used to calculate the female index (FI):

FI = (number of females on a given genotype/average number of females on the susceptible control) × 100 (Niblack et al., 2009).

All plants were maintained in the growth chamber (Percival, Perry, IA, USA) under controlled conditions at 27 °C, 16 h light/8 h dark, and 50% relative humidity throughout the assay.

For nematode preparation, cysts were collected from stock roots that had been maintained in the same greenhouse. Briefly, SCN cysts were harvested by massaging the stock roots in water and sieving the solution through nested 850- and 250-µm test sieves (Fisher Scientific, Suwanee, GA, USA). The collected cysts were crushed with a rubber stopper in a 250-µm sieve, and the released eggs were collected in a 25-µm mesh sieve. The eggs were then purified by sucrose flotation (Matthews et al., 2003) with some modifications. For nematode hatching, purified eggs were placed on wet paper tissue in a plastic tray with an appropriate level of water, covered with aluminum foil, and maintained at 27 °C for 3 days. Hatched second-stage juvenile nematodes (J2) were collected and suspended in 0.09% liquid agarose at a final concentration of 1,800 J2 ml⁻¹. For inoculation, 1 ml of J2 inoculum was added to each root as treatment, and seedlings inoculated with 0.09% agarose were used as non-infected controls.

Three days post-inoculation (dpi), three roots from randomly selected seedlings were stained with acid fuchsin to validate successful inoculation and investigate nematode development in the roots as previously described (Bybd et al., 1983, Klink et al., 2007b).

For transcriptome sequencing and LC-MS analyses, root tissues were separately prepared and sampled at 3, 5, and 8 dpi. The rationale for choosing these three time points was based on: 1) our previous study showing that SCN development began to differ between WsR and WsS roots at approximately 5 dpi, with the difference becoming more apparent at 8 dpi (Zhang et al., 2017c); and 2) previous studies of SCN infection reporting differences in syncytium and nematode development between resistant and susceptible roots at 5–8 dpi (Kandoth et al., 2011, Klink et al., 2007b).

For RNA-seq (Zhang et al., 2017c, Zhang et al., 2017b), root tissues from four individual plants were pooled at each sampling time for each biological replicate, and three biological replicates were collected for each condition. For each biological replicate, root tissues collected at 3, 5, and 8 dpi were subsequently pooled before RNA extraction and library construction. Thus, the RNA-seq dataset represents an integrated transcriptional response across the 3–8 dpi infection window rather than independent time-resolved transcriptomes.

For LC-MS analysis, roots from 3–5 plants per condition were collected, weighed, and analysed individually at each time point. All samples were immediately frozen in liquid nitrogen and stored at −80 °C.

### RNA extraction, transcriptome sequencing, and data analyses

RNA was isolated using an RNeasy Plant Mini Kit (Qiagen, Hilden, Germany) following the manufacturer’s instructions and quantified using a NanoDrop 2000 (Thermo Fisher Scientific, Waltham, MA, USA). RNA integrity, purity, and concentration were assessed using an Agilent 2100 Bioanalyzer with an RNA 6000 Nano Chip (Agilent Technologies, Santa Clara, CA, USA). Prior to library construction, total RNA was treated with RNase-free DNase I (New England Biolabs, Ipswich, MA, USA) to remove contaminating genomic DNA.

Messenger RNA (mRNA) was purified using the NEBNext Poly(A) mRNA Magnetic Isolation Module (New England BioLabs, Ipswich, MA, USA) following the manufacturer’s directions. Complementary DNA (cDNA) libraries for Illumina sequencing were constructed using the NEBNext Ultra Directional RNA Library Prep Kit and NEBNext Multiplex Oligos for Illumina (New England BioLabs) according to the manufacturer’s protocol, as previously described (Zhang et al., 2017c). The amplified library fragments were purified and checked for quality and final concentration using an Agilent 2200 TapeStation (Agilent Technologies, Santa Clara, CA, USA). The final quantified libraries were pooled in equimolar amounts for sequencing on an Illumina HiSeq 2500 with a read length of 125 bp and v4 sequencing chemistry (Illumina, San Diego, CA, USA).

Enrichment analyses for GO terms and KEGG pathways were performed as previously described (Zhang et al., 2017c). For construction of the functional association network, functional associations among selected genes were retrieved from SoyNet (Kim et al., 2017) and visualized using Cytoscape (Shannon et al., 2003).

### Quantitative real-time PCR analysis

RNA extraction from root tissues for qPCR and genomic DNA removal using DNase I were conducted using the same protocols described above. Reverse transcription reactions were performed using the RevertAid First Strand cDNA Synthesis Kit (Thermo Fisher Scientific, Waltham, MA, USA) following the manufacturer’s instructions. Quantitative real-time PCR (qPCR) was performed using PerfeCTa™ SYBR® Green FastMix™ (Quanta Biosciences, USA) on an ABI 7500 Fast real-time PCR system (Applied Biosystems, USA).

Three biological replicates per sample were used for qPCR, and each reaction was repeated twice as technical replicates. The soybean ubiquitin 3 gene (GmUBI-3, accession D28213) was used as an endogenous control (Zhang et al., 2017c, Kandoth et al., 2011). All primers used in this study are listed in Table S2.

### Metabolite extraction, data processing, and data analysis

Metabolites were extracted from each root tissue, and the resulting data were processed as previously described (Strauch et al., 2015). Briefly, root tissues were extracted with 50% (v/v) methanol in a 60 °C water bath for 30 min. A tissue-to-solvent ratio (w/v) of 1:10 was used consistently across all samples. Extracts were filtered using a 0.2-µm filter prior to LC-MS profiling on a G6530A Q-TOF LC/MS (Agilent Technologies, USA). Peaks consistently detected in at least three biological replicates within each group at each time point were used for downstream analysis.

The identities of selected metabolites were confirmed using authentic standards, including daidzein, daidzin, genistein, genistin, formononetin, SA (2-hydroxybenzoic acid), 3-hydroxybenzoic acid, 4-hydroxybenzoic acid, 2,3-dihydroxybenzoic acid, and 4-hydroxybenzaldehyde. Remaining metabolite features were putatively annotated based on mass matching against KEGG (Kanehisa et al., 2016), the Plant Metabolic Network (PMN)/SoyCyc database, and an in-house database.

Analysis of metabolomics data was performed using MetaboAnalyst (Xia et al., 2015). Briefly, peak-area data were cube-root transformed and Pareto scaled prior to statistical analysis (Alonso et al., 2015). One-way ANOVA with Fisher’s LSD post hoc analysis (FDR < 5%) was performed on normalized data to determine differentially abundant metabolites among four groups (WsR_T, WsR_C, WsS_T, and WsS_C) at each time point.

For data visualization, PCA was conducted in R using princomp using all detected high-confidence metabolite features. Heat maps illustrating metabolite abundance patterns were generated using JMP Pro 13 (SAS Institute Inc., Cary, NC, USA). Biological databases, including KEGG, MetaCyc, and PMN/SoyCyc, were used as references to map metabolites to their corresponding pathways.

### Nematicidal activity assay

Determination of the nematicidal activity of phenolic compounds was conducted as previously described (Nguyen et al., 2013). Briefly, approximately 400 freshly hatched J2s in 900 µl of water were added to each well of a 24-well Microtest™ tissue culture plate, and 100 µl of compound test solution was added at concentrations of 1.25, 2.5, 5, and 10 mg ml⁻¹, resulting in final concentrations of 0.125, 0.25, 0.5, and 1.0 mg ml⁻¹, respectively. These concentrations were used in a previous study examining the nematicidal activity of 3,4-dihydroxybenzoic acid against root-knot nematodes (Nguyen et al., 2013). Control samples received 100 µl of water. Three biological replicates per concentration were used.

The plate was covered with the original solid lid and wrapped with Parafilm®, and samples were kept at 27 °C in an incubator. After 24 h of incubation, nematodes were considered paralyzed when no movement was observed during 2 seconds of observation after prodding with a fine needle (Cayrol et al., 1989). To determine whether paralysis was reversible, paralyzed nematodes were transferred to distilled water. Nematodes that showed no recovery of movement after 48 h were considered dead. All J2s were counted using a Leica M165 FC stereomicroscope (×40) to determine mortality. J2 mortality was calculated as the percentage of dead J2s relative to the total number of J2s examined. The compounds 2,3-dihydroxybenzoic acid and 4-hydroxybenzaldehyde were obtained from Sigma-Aldrich, USA.

## Supplementary figure legends

**Supplementary Fig. 1 Multidimensional scaling and comparison of DEGs between genotypes.**

**a** Multidimensional scaling (MDS) plots of RNA-Seq samples from WsR and WsS following inoculation with SCN races 2 and 5 and their corresponding controls. Biological replicates cluster according to experimental condition. **b** Venn diagrams showing overlap among up- and downregulated DEGs identified in WsR and WsS in response to races 2 and 5.

**Supplementary Fig. 2 Expression patterns and functional enrichment of DEGs.**

**a** Heat maps showing expression patterns of DEGs induced in WsR by SCN races 5 and 2 across the four genotype-by-race conditions. **b** GO enrichment analysis of downregulated rDEGs in response to race 2. **c** GO enrichment analysis of downregulated rDEGs in response to race 5. **d** GO and KEGG enrichment analyses of 259 upregulated cDEGs. P, F, and C denote biological process, molecular function, and cellular component, respectively.

**Supplementary Fig. 3 Comparative expression analysis of selected DEGs.**

**a** Expression patterns of CAMTA4 (Glyma.11G251900) and SAMT1 (Glyma.02G054200) across the four genotype-by-race conditions. **b** Time-course expression analysis of GmICS1 (Glyma.03G070600) and GmICS2 (Glyma.01G104100) at 3, 5, and 8 dpi across the four genotype-by-race conditions. R2 and R5 denote SCN races 2 and 5, respectively.

