## Supplemental Figures for "Phenolic Chemical Defense Contributes to Resistance against Multiple Soybean Cyst Nematode Populations in Wild Soybean"

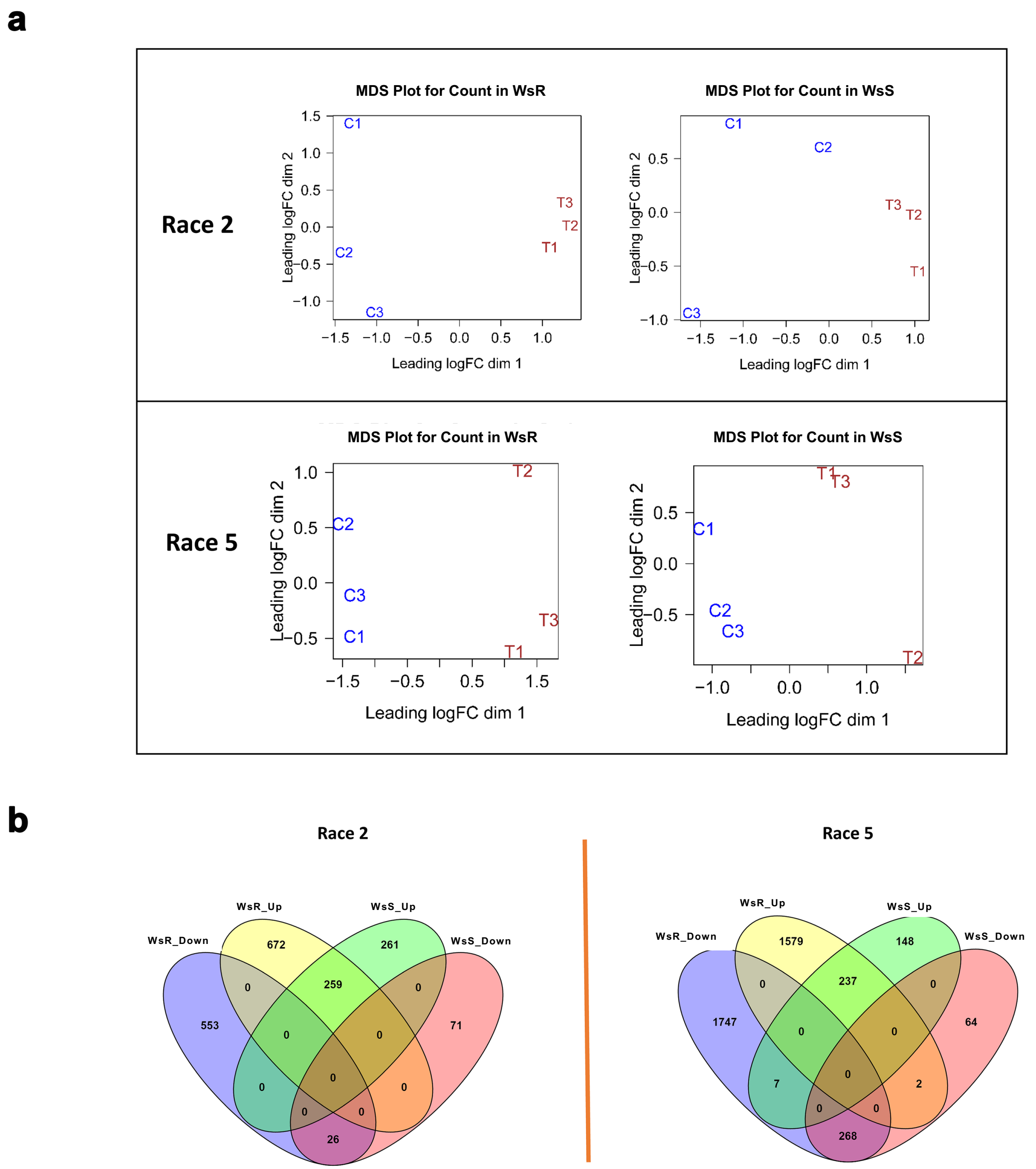
a
Race 2
MDS Plot for Count in WsR
MDS Plot for Count in WsS
Leading logFC dim 1
Leading logFC dim 2
C1
C2
C3
T1
T2
T3
Race 5
MDS Plot for Count in WsR
MDS Plot for Count in WsS
Leading logFC dim 1
Leading logFC dim 2
b
Race 2
WsR_Down
WsR_Up
WsS_Up
WsS_Down
672
261
553
259
71
26
Race 5
WsR_Down
WsR_Up
WsS_Up
WsS_Down
1579
148
1747
237
64
7
2
268

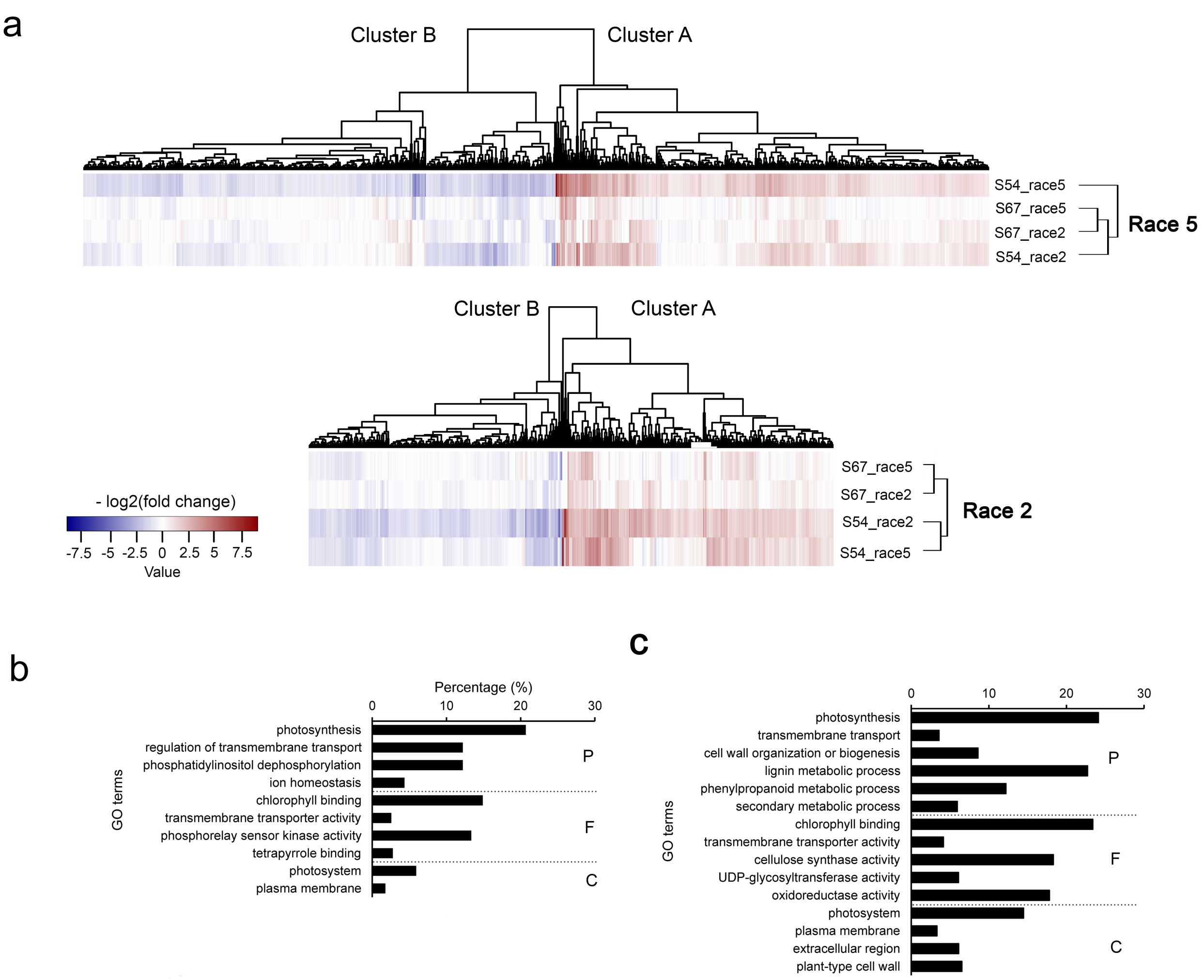


d

**GO enrichment for Cm259Up**

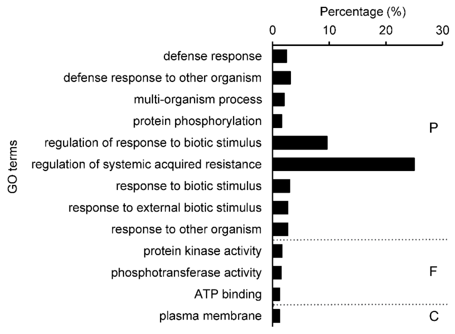


**KEGG enrichment for Cm259Up**

| Term | Database | ID | *P*-Value |
|---|---|---|---|
| Plant-pathogen interaction | KEGG PATHWAY | gmx04626 | 0.027329845 |
| Monoterpenoid biosynthesis | KEGG PATHWAY | gmx00902 | 0.040939836 |

P: biological process
F: molecular function
C: cellular componet

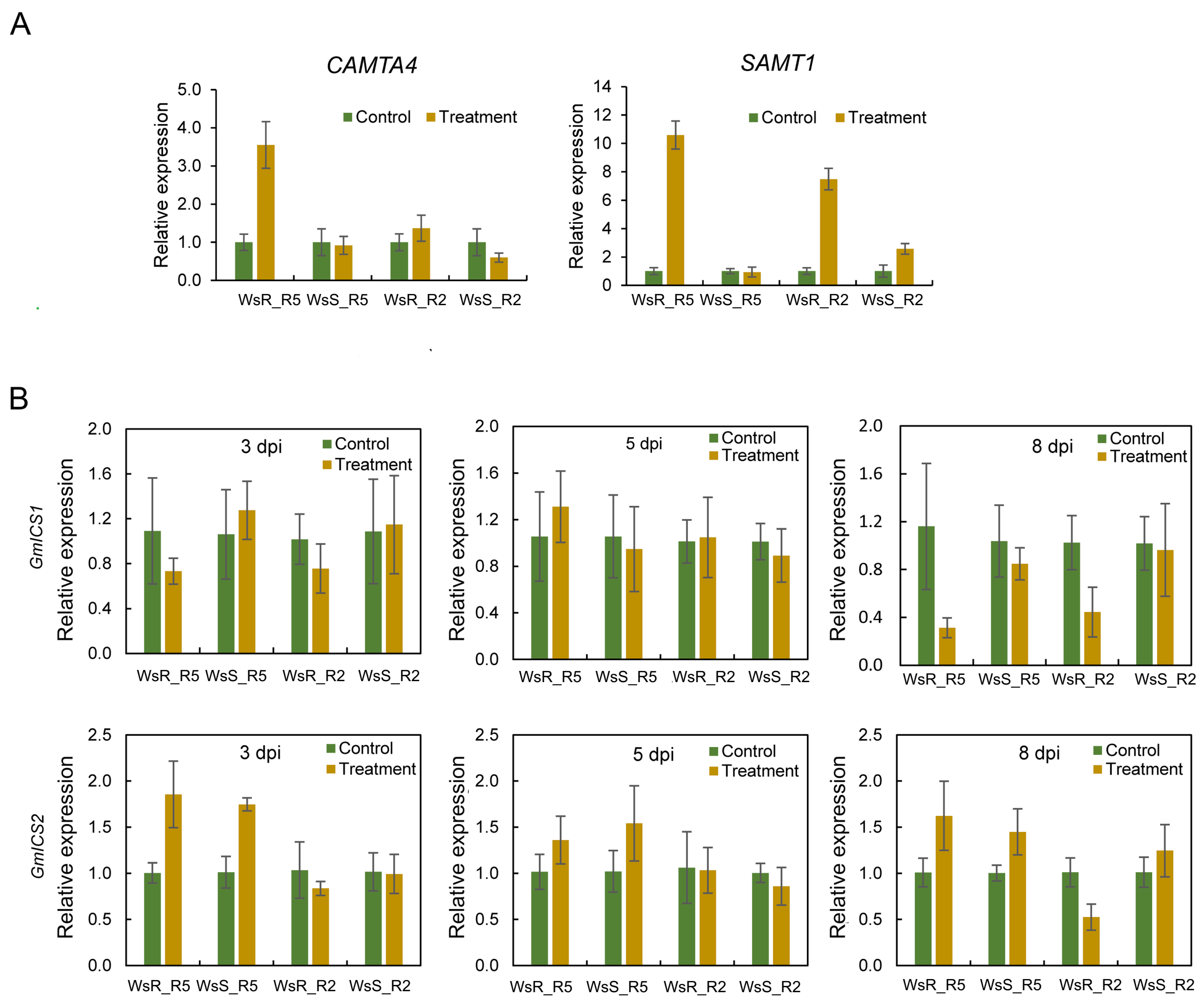
A
CAMTA4
SAMT1
Relative expression
Control
Treatment
WsR_R5
WsS_R5
WsR_R2
WsS_R2
B
GmICS1
GmICS2
3 dpi
5 dpi
8 dpi
